# Targeting DNA Polymerase Epsilon Leads to Tumor Clearance and Activation of an NF-κB-mediated inflammatory response in Triple Negative Breast Cancer

**DOI:** 10.1101/2025.09.18.677147

**Authors:** Elizabeth F. Sher, Kenji M. Fujihara, Anthony Tao, Diana Erenburg, Vladislav O. Sviderskiy, Hannan Mir, Triantafyllia R. Karakousi, Cynthia Loomis, Jiehui Deng, Kwok-Kin Wong, Richard Possemato

## Abstract

Breast cancer remains the second leading cause of cancer-related mortality among women, with triple-negative breast cancer (TNBC) exhibiting a particularly poor five-year prognosis^1^. Here, we demonstrate that among genetic and pharmacological perturbations targeting DNA replication, suppression of DNA polymerase epsilon (POLE) in TNBC, induces a potent, TNBC-specific gene expression signature enriched in inflammatory cytokines that are transcriptional targets of NF-κB. TNBC cells exhibit markedly higher levels of DNA damage and canonical NF-κB activation compared to luminal breast cancer cells. Notably, NF-κB activation in this context depends on the canonical component RELA, but not the non-canonical component RELB. Mechanistically, ATM, STING, and RIG-I each contribute to NF-κB activation following POLE suppression. In vivo, POLE suppression in a murine TNBC model leads to cancer cell-intrinsic elimination of tumor burden and increased immune cell infiltration. Together, these findings support a model in which replication stress from POLE inhibition triggers robust NF-κB–mediated inflammation and immune microenvironment remodeling in TNBC and can independently trigger tumor eradication. These results suggest a potential therapeutic avenue for targeting POLE in TNBC.

## INTRODUCTION

Breast cancer is the second leading cause of cancer-related mortality in US women^2^. Breast cancer can be classified by estrogen receptor (ER) or progesterone receptor (PR) expression and amplification of HER2, while breast tumors lacking these markers are designated as triple-negative (TNBC). Expression profiling has yielded molecular classifications and further divided TNBC into distinct subtypes, some of which have immune activated signatures that are therapeutically relevant^3,4^. ER+/PR+ luminal breast cancer (LUBC) and HER2-amplified subtypes are successfully targeted by hormone therapy and HER2 blocking antibodies, respectively. In contrast, no similarly effective targeted therapies are currently available for TNBC. TNBC displays frequent relapse, poor prognosis despite relative chemosensitivity, and disproportionate mortality^5^. Immune checkpoint inhibitors (ICI) have met with some success in TNBC independent of mutational burden^4,6–8^. However, patients harboring TNBC tumors that present with high levels of immune infiltration have markedly better prognosis, raising the hope that approaches that trigger immune cell infiltration may promote anti-cancer responses alone or in combination treatments^9,10^.

NF-κB is a key transcription factor that modulates the tumor microenvironment (TME) by influencing immune and stromal cell populations, including macrophages, dendritic cells, myeloid-derived suppressor cells, neutrophils, NK cells, NKT cells, T and B lymphocytes, cancer-associated fibroblasts, and endothelial cells^11^. NF-κB also drives the senescence-associated secretory phenotype, which promotes innate immune cell recruitment to the TME^12^. In TNBC, chronically elevated NF-κB activity is associated with poor prognosis, and pathway inhibition induces tumor control in some contexts^13^. In contrast, acute activation of NF-κB may alter the TME in ways that are immunostimulatory^14–16^.

DNA damage is a potent inducer of NF-κB activation, with ataxia-telangiectasia mutated (ATM) kinase playing a central role in this process^17–19^. Canonical NF-κB signaling is activated through the IKK complex (IKKα, IKKβ, and IKKγ/NEMO), which regulates the cytoplasmic sequestration and release of the p50/p65 heterodimer via IκBα^20,21^. In contrast, the non-canonical NF-κB pathway involves the de novo synthesis of NF-κB-inducing kinase (NIK) and culminates in p52/RelB-mediated transcriptional responses^22,23^. Genotoxic stress triggers NF-κB activity, with dual roles in cell survival and apoptosis. DNA damage-induced NF-κB activation can protect cells from apoptosis^24^, yet prolonged genotoxic stress may promote cell death via TNF and IL-8 secretion, forming a potential autocrine feedback loop^25^.

In addition to DNA damage sensing through the nucleus-to-cytoplasm axis, NF-κB can be activated via pattern recognition receptors (PRRs) such as TLRs^26^, MAVS^27^, STING^28^, and NOD-like receptors^29^, as well as antigen receptors on T and B cells^30,31^. Double-stranded DNA (dsDNA) breaks act as danger-associated molecular patterns (DAMPs), initiating rapid innate immune responses via NF-κB activation. Emerging evidence implicates cGAS-STING, RIG-I, and p38 MAPK in relaying genotoxic signals to NF-κB^32–35^. Indeed, replication stress and subsequent release of dsDNA upon treatment with PARP inhibitors can modulate expression of programmed cell death 1 ligand 1 (PDL1), whose targeting by immunotherapy was recently shown to improve overall survival in TNBC.^36,37^ As such, understanding mechanisms of inflammatory signaling pathway activation by replication stress can be leveraged to improve future TNBC therapies.

Three polymerase complexes duplicate the mammalian genome: DNA polymerases alpha (POL α), delta (POL δ), and epsilon (POL ε), nucleated by catalytic subunits POLA1, POLD1, and POLE, respectively.^38^ POL δ primarily performs lagging strand synthesis, POL ε synthesizes the leading strand, and POL α extends from the RNA primer and is required for synthesis of both strands.^39^ There are no anti-cancer drugs that directly target DNA replicative polymerases, and available inhibitors of replication (aphidicolin) or dNTP synthesis (hydroxyurea, gemcitabine, 5-fluorouracil) are non-selective, blocking replicative polymerases as well as a cadre of polymerases involved in DNA repair.

We have previously shown that TNBC is especially sensitive to partial suppression of POLE^40^. However, it is unclear whether the response to POLE inhibition is distinct from other mechanisms of replication targeting, including those that are non-specific or induce DNA damage through other mechanisms. Here, we show that suppression of Pol ε in TNBC cells, but not LUBC cells, leads to a distinct transcriptional signature characterized by an increase in inflammatory cytokine transcripts known to be targets of the NF-κB transcription factor. TNBC cells display markedly higher levels of DNA damage and canonical NF-κB activation compared to LUBC cells upon POLE inhibition. Notably, NF-κB activation in this context is dependent on the canonical pathway component RELA, but not on the non-canonical component RELB. Mechanistically, ATM, STING, and RIG-I each contribute to NF-κB activation following POLE suppression. In murine models of TNBC, POLE inhibition leads to tumor regression and alters immune infiltration, indicating that POLE can be therapeutically targeted.

## RESULTS

### Inhibition of leading strand replication in TNBC activates a transcriptional response distinct from DNA-damaging agents or inhibition of lagging strand replication

Our prior work demonstrated that TNBC cell lines exhibit a selective requirement for POLE compared to LUBC or non-transformed mammary epithelial cells.^40^ Analysis of large-scale genetic screening data validate these findings, as POLE is the top selective liability in TNBC (Fig. 1A, Supplementary Table 1). This analysis successfully identifies both clinically effective targets in LUBC (Cyclin D/CDK4/6 and estrogen receptor signaling) as non-TNBC-selective. Indeed, compared to other cancer subtypes, TNBC exhibits the highest dependence on POLE (Fig. 1B, Supplementary Table 1). In contrast, no selective requirement is observed for POLA1, POLD1, and ribonucleotide reductase (RRM2), responsible for dNTP generation (Fig. 1A), indicating that leading strand replication is a specific vulnerability in TNBC.

**FIGURE 1:**
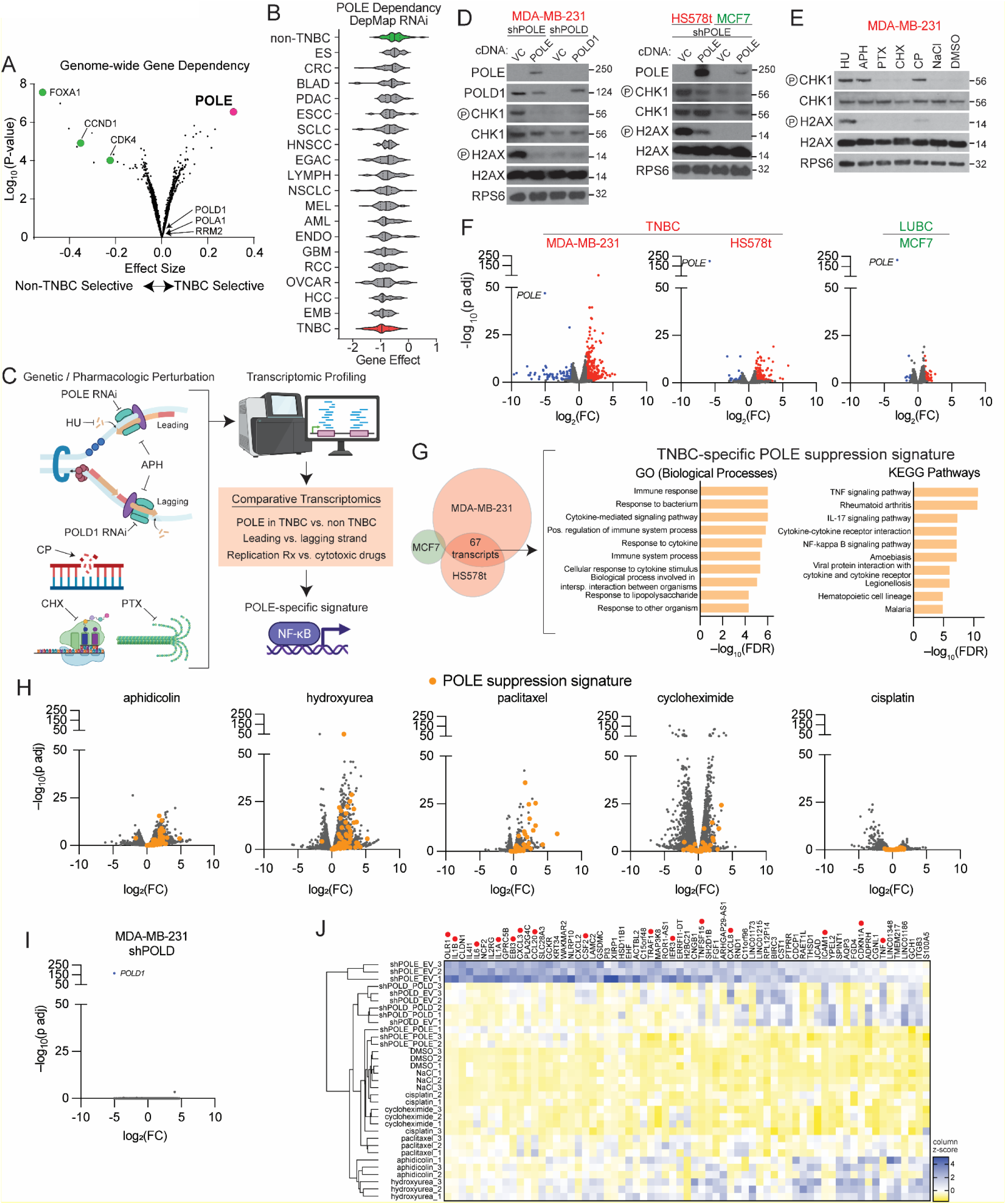
Inhibition of leading strand replication in TNBC activates a transcriptional response distinct from DNA-damaging agents or inhibition of lagging strand replication. **A,** Genome-wide RNAi screening gene effect data (DepMap) comparing 29 TNBC and 46 non-TNBC lines. POLE is the most selectively required gene in TNBC (pink circle). The most selectively required genes in non-TNBC lines include ER transcription factor FOXA1 and palbociclib targets CDK4 and cyclin D1 (green circles). **B,** POLE dependency (DepMap gene effect) as in A for each indicated tumor subtype. **C,** Diagram of comparative transcriptomic profiling for indicated chemotherapeutic and genetic perturbations. HU (Hydroxyurea), Aph (Aphidicolin), CP (Cisplatin), CHX (Cycloheximide), PTX (Paclitaxel). **D,** Immunoblots for indicated proteins of lysates from indicated cell lines transduced with indicated shRNAs and expressing indicated cDNAs or vector control (VC), 5d. **E,** Immunoblot of lysates from MDA-MB-231 cells with HU (2mM), Aph (2μg/mL), CP (12μM), CHX (2μM), PTX (4nM) or vehicle (NaCl or DMSO), 24 hrs. **F,** Volcano plot showing genes differentially increased (red) or decreased (blue) in cell lines from *D* using an adjusted p-value cutoff of 0.05 and log_2_FC cutoff of ±1. **G,** Venn diagram indicated overlap of differentially expressed genes from *F*, the corresponding gene sets enriched in the commonly upregulated 67 genes, and their degree of statistical significance. **H,** Volcano plot showing genes differentially expressed in cell lines from E, with those 67 genes identified in F colored orange. **I,** Volcano plot as in *F* for POLD suppression. **J,** Heatmap showing differential gene expression of the 67 genes identified in *F* across all samples analyzed grouped by hierarchical clustering. Genes known to be NF-κB targets indicated by red circles. Immunoblots are representative of n=3 independent replicates.

To understand how inhibition of POLE might be distinct from other methods of targeting DNA replication, we used genetic and pharmacological methods to perturb replication in TNBC and LUBC cell lines (Fig. 1C). Two TNBC cell lines (MDA-MB-231 and HS578T) and one luminal breast cancer (MCF7) cell line were engineered to express either an empty vector (MXS EV) or an shRNA-resistant POLE or POLD cDNA (MXS POLE and MXS POLD), followed by shRNA-mediated suppression of endogenous POLE or POLD (Fig 1D). As we had observed previously,^40^ suppression of POLE in TNBC cell lines caused marked DNA damage (measured by H2AX phosphorylation at S139 [pH2AX]) and replication fork stalling (measured by phosphorylation of Chk1 at S345 [pCHK1]). In contrast, LUBC cell line MCF7 exhibited no DNA damage or replication fork stalling in response to partial POLE suppression (Fig. 1D). We also treated MDA-MB-231 cells with a panel of replication stress-inducing agents and chemotherapeutics for 24 hours. These included aphidicolin (APH; 2 µg/mL), which inhibits DNA polymerases α, δ, and ε; hydroxyurea (HU; 2 mM), which depletes nucleotide pools by inhibiting ribonucleotide reductase; paclitaxel (PTX; 4 nM), which induces mitotic stress that indirectly perturbs DNA replication by arresting cells in metaphase; cycloheximide (CHX; 2 µM), which indirectly impairs replication by blocking protein synthesis and depleting essential replication factors; and cisplatin (CP; 12 µM), which forms DNA crosslinks, leading to replication fork collapse and DNA damage. The doses of these genotoxic agents were selected to induce cell death within 48 hours of treatment, and samples were harvested at 24 hours, prior to overt cell death. Consistent with their mechanism of action, treatment with HU, APH and CP led to replication fork stalling and DNA damage, and this phenotype was most pronounced in HU-treated samples (Fig 1E).

To understand whether POLE inhibition and treatment with genotoxic agents in these breast cancer models leads to corresponding changes in gene transcription, we performed an unbiased comparative transcriptomic analysis (Fig. 1C, Supplementary Table 2). POLE suppression in TNBC cell lines led to the selective upregulation of 67 transcripts compared to LUBC, forming a TNBC-specific POLE gene signature (Figure 1F-G, Supplementary Table 3). To explore the functional relevance of this signature, we performed pathway enrichment analysis using STRING^41^ and identified the top 10 enriched biological processes. The analysis revealed significant upregulation of immune and inflammatory pathways, including IL-17 signaling, cytokine-mediated signaling, NF-κB signaling, and other pro-inflammatory processes (Figure 1G).

We then asked whether inhibition of leading strand replication induces a transcriptomic signature in TNBC similar to inhibition of lagging strand replication and treatment with DNA-targeting genotoxic and chemotherapeutic agents (Fig. 1E). Among all agents tested, aphidicolin and hydroxyurea induced transcriptional responses most similar to POLE suppression, consistent with their ability to impair DNA replication and cause replication fork stalling (Figure 1H-J, Supplementary Table 1). Paclitaxel, a frontline chemotherapeutic for TNBC known to enhance immunotherapy efficacy^42^, also modestly upregulated POLE-associated transcripts, indicating that mitotic stress alone is insufficient to trigger inflammatory signaling. Cycloheximide induced widespread transcriptional changes that shared little overlap with the POLE suppression signature (Figure 1H, J). Cisplatin—a widely used frontline chemotherapeutic^43^ and potent DNA-damaging agent—also showed minimal overlap with the POLE suppression signature and induced few POLE-associated transcripts (Figure 1H, J). The magnitude and specificity of inflammatory gene induction were more pronounced in POLE-suppressed cells than all genotoxic treatments (Figure 1J), suggesting that POLE suppression activates a distinct inflammatory transcriptional response. In particular, a number of inflammatory cytokines (IL-6, IL-1α, IL-1β, CXCL2, and CXCL3) are highly upregulated in TNBC cells upon POLE suppression. Fourteen of these 67 genes are commonly targeted by the NF-κB family of transcription factors, indicating that activation of this pathway is a major feature of POLE inhibition (Figure 1J, red dots, Supplementary Table 3).

### POLE and POLD1 exhibit distinct threshold requirements for cell viability and subtype-specific effects on inflammatory gene expression

Suppression of POLD1 resulted in minimal transcriptional changes, despite robust suppression of POLD1 protein levels (Figure 1I). To understand whether this observation reflected a different threshold for POLE and POLD1 requirement, we expressed POLE and POLD1 cDNAs under doxycycline (DOX) control, followed by CRISPR-Cas9-mediated deletion of the corresponding endogenous gene (DOX-OFF POLE, DOX-OFF POLD1), or partially suppressed these genes with an shRNA, in MDA-MB-231 and MCF7 cells. DOX addition led to near-complete suppression of the corresponding protein, and a similar induction of DNA damage and replication fork stalling in both TNBC and LUBC models (Figure 2A-B), whereas only RNAi-mediated suppression of POLE in TNBC exhibited a substantial increase in DNA damage (Figure 2C). Accordingly, RNAi-mediated suppression of POLE reduced cell viability selectively in MDA-MB-231 cells, whereas near-complete POLD1 and POLE suppression strongly reduced cell viability in both TNBC and LUBC models (Figure 2D-E). Consistent with induction of DNA damage in MDA-MB-231 cells, near-complete POLE and POLD1 suppression transcriptionally activates a panel of NF-κB-associated cytokines (IL-6, IL-1α, IL-1β, CSF3) identified in our transcriptomic analysis (Fig. 1J). However, near-complete suppression of either POLE or POLD1 does not substantially activate this response in MCF7 cells despite the presence of robust H2AX phosphorylation (Figure 2F), demonstrating that TNBC is inherently more capable of activating an NF-κB-associated gene expression profile in response to DNA damage. These data further demonstrate that POLD1 expression is in substantial excess of its requirement for cell viability and genomic integrity, whereas POLE expression is limiting, particularly in TNBC models. Thus, POLE is a selective liability in TNBC and partial inhibition of POLE potently activates a specific transcriptional response that includes NF-κB activation.

**FIGURE 2:**
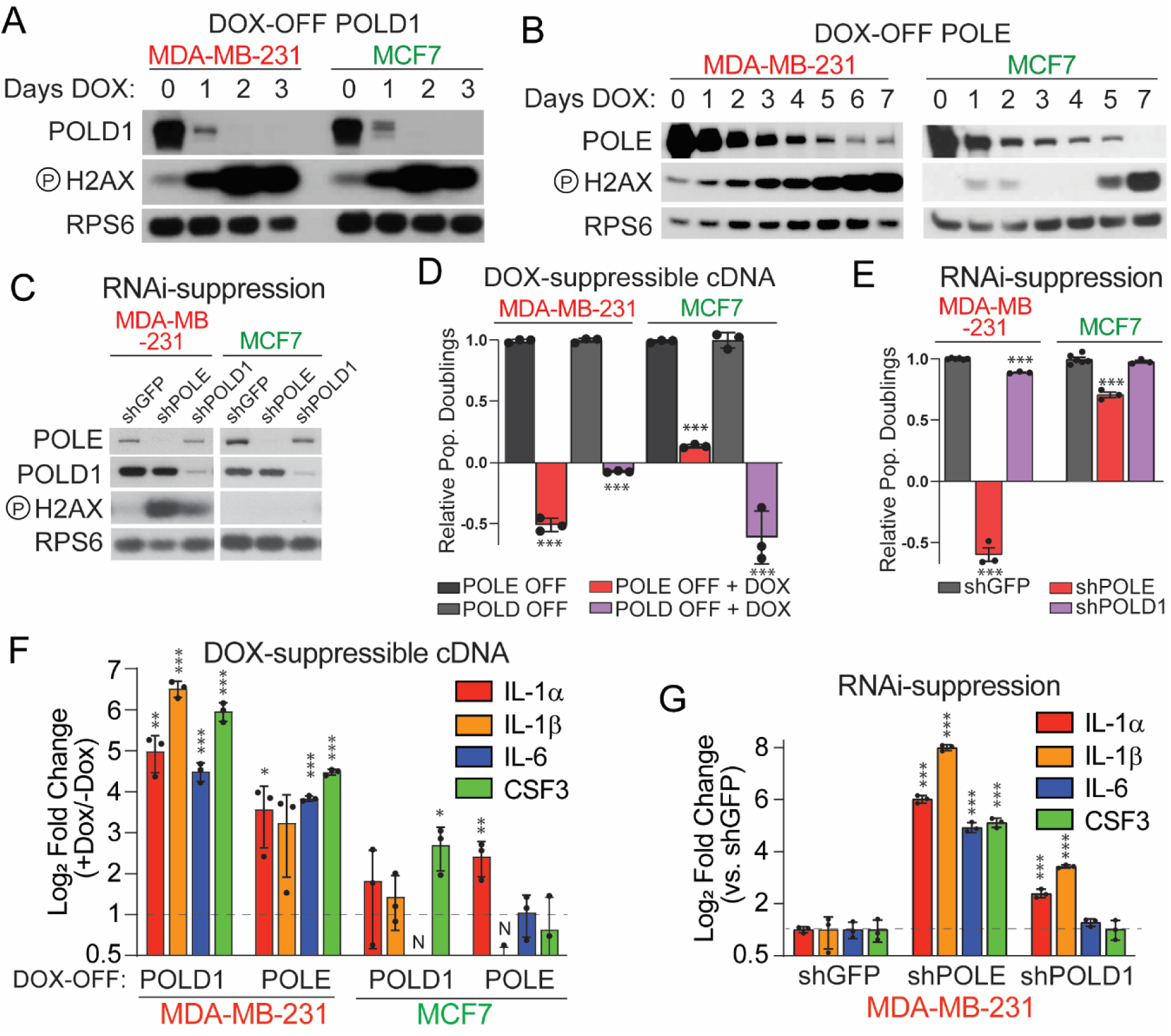
POLE and POLD1 exhibit distinct threshold requirements for cell viability and subtype-specific effects on inflammatory gene expression. **A-B,** Immunoblots for indicated proteins of lysates from indicated cell lines expressing a doxycycline (DOX)-repressible cDNA or POLD1 or POLE and CRISPR/Cas9-mediated deletion of endogenous POLD1 (DOX-OFF POLD1) or POLE (DOX-OFF POLE) after the indicated duration of DOX treatment (0.1μg/mL). **C,** Immunoblots for indicated proteins of lysates from indicated cell lines expressing shRNAs targeting POLD1 (shPOLD), POLE (shPOLE) or control (shGFP). **D,** Population doublings of cells in *A-B* after treatment with DOX for 3d, 5d growth. **E,** Population doublings of cells in *C* 5d after shRNA transduction, 5d growth. **F,** Fold change in mRNA levels (log_2_) of indicated genes upon DOX addition (5d) of cell lines from *A-B*. **G,** Fold change in mRNA levels (log_2_) of indicated genes upon shRNA transduction (5d) of cell lines from *C*. Immunoblots are representative of n=3 independent replicates. Data in ***D-G*** report the mean and standard deviation of n=3 biological replicates, *** p<0.001, ** p<0.01, *p<0.05.

### POLE inhibition induces RELA-dependent NF-κB activation and inflammatory cytokine release

Given the robust increase in cytokine transcripts targeted by NF-κB upon POLE suppression, we assessed NF-κB pathway signaling and activity. Canonical NF-κB signaling is regulated by differential phosphorylation of inhibitor of kappa-B-alpha (IκBα), resulting in its selective degradation and subsequent activation of RELA. Accordingly, POLE suppression increases phosphorylation of RELA and decreases levels IκBα (Figure 3A). We then evaluated NF-κB activation using a reporter containing multimerized κB-binding sites adjacent to a minimal promoter and firefly luciferase. We observed a significant increase in NF-κB activity upon POLE suppression in TNBC cell line MDA-MB-231, similar to TNFα treatment (Figure 3B). Expression of a phosphorylation- and degradation-resistant IκBα-SS32/36AA super repressor (IκBα-SR), increases total IκBα protein and baseline RELA phosphorylation (Figure 3A) and prevents activation of the NF-κB luciferase reporter upon POLE suppression (Figure 3C).

**FIGURE 3:**
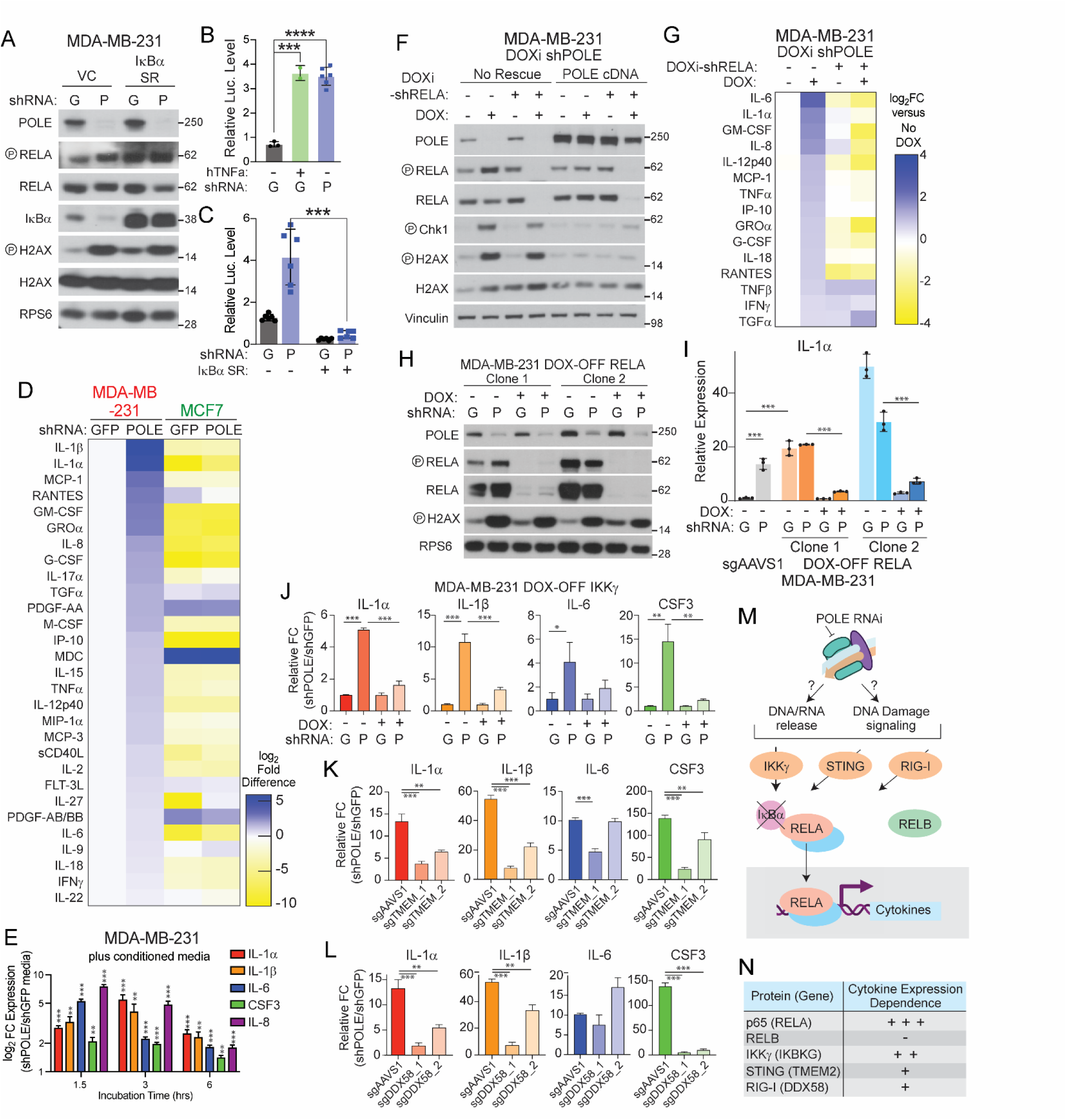
POLE inhibition induces RELA-dependent NF-κB activation and inflammatory cytokine release. **A-B,** Immunoblots for indicated proteins of lysates from MDA-MB-231 cells expressing a control vector (VC) or NF-kB super-repressor (IκBα SR) and shRNAs targeting POLE (P) or a control (shGFP, G). **B-C**, Luminescence of cells from ***A***, with indicated TNFα treatment (hTNFa, 20ng/ml for 20 minutes). **D**, Fold change (log_2_) in the level of indicated cytokines and chemokines in cell culture media of indicated cell lines transduced with indicated shRNAs (5d), relative to MDA-MB-231 cells transduced with shGFP, n=2. **E**, Fold change (log_2_) in the mRNA expression of the indicated genes in MDA-MB-231 cells exposed to media conditioned for 5d by MDA-MB-231 cells transduced with shPOLE versus those transduced with shGFP for the indicated durations (hrs). **F**, Immunoblots for indicated proteins of lysates from MDA-MB-231 cells expressing a DOX-inducible shRNA targeting POLE and a DOX-inducible shRNA targeting RELA or an shRNA-resistant POLE cDNA, as indicated. DOX addition as indicated (100ng/mL, 5d). **G**, Fold change (log_2_) in the level of indicated cytokines and chemokines in cell culture media from the indicated cell lines from ***F***, n=2. **H,** Immunoblots for indicated proteins of lysates from MDA-MB-231 cells expressing a DOX-repressible RELA cDNA and endogenous *RELA* disruption. Transduction with shRNAs targeting POLE and DOX addition (100ng/mL) as indicated (5d). **I,** mRNA level of IL-1α in cell lines from ***H*** or those expressing a control sgRNA (sgAAVS1), relative to sgAAVS1 clones transduced with shGFP (5d). **J,** mRNA level of indicated genes from MDA-MB-231 cells expressing a DOX-repressible IKKγ cDNA and endogenous *IKKγ* disruption, relative to shGFP transduced cells. Transduction with shRNAs targeting POLE and DOX addition (100ng/mL) as indicated (5d). **K-L,** mRNA level of indicated genes from MDA-MB-231 knockout clones generated using the indicated sgRNAs and transduced with shPOLE or shGFP (5d), relative to shGFP transduced cells in each condition. **M,** Schematic of interrogated signaling pathways downstream of replication stress. **N,** Summary of the contribution of interrogated signaling pathways to expression of NF-κB-regulated cytokines. Immunoblots are representative of n=3 independent replicates. Data in ***B***, ***C***, ***D***, ***E***, ***G***, ***I-L*** report the mean and standard deviation of n=3 biological replicates. *** p<0.001, ** p<0.01, *p<0.05.

We then considered the impact of POLE suppression on cytokine secretion. POLE suppression induces robust increases in the extracellular levels of numerous cytokines and chemokines in MDA-MB-231 cells, which are unaltered in MCF7 cells (Figure 3D). We validated a subset of these in an expanded cell line panel (Supplementary Figure 1A). To determine whether cytokine release contributes to an autocrine signaling loop, we treated naive MDA-MB-231 cells with conditioned media from POLE-suppressed cells. This treatment rapidly induced expression of several NF-κB-associated cytokine transcripts, supporting a feed-forward loop of inflammatory amplification (Figure 3E).

To further test whether canonical NF-κB pathway signaling is necessary for this transcriptional response, we generated cells stably expressing a DOX-inducible POLE shRNA (DOXi-shPOLE) with or without a DOX-inducible RELA shRNA (DOXi-shRELA) and POLE cDNA rescue. We confirmed that DOX addition resulted in the expected reduction of POLE and RELA, and that the effects of POLE suppression on DNA damage and replication fork stalling could be rescued by POLE cDNA expression (Figure 3F). H2AX and CHK1 phosphorylation are not impacted by RELA suppression, indicating that NF-κB activation is not required for replication fork stalling and DNA damage upon POLE suppression (Figure 3F). In this model, the secretion of 13 cytokines was induced by greater than 50% upon DOX-induced POLE suppression, and RELA inhibition blocked this increase in 12 of these cytokines (Figure 3G). We additionally generated a DOX-OFF RELA system in which DOX addition induces a transition between a RELA-overexpressed and a RELA knockout state by transduction of a RELA cDNA under DOX control followed by CRISPR-Cas9 mediated KO of endogenous *RELA* (Figure 3H). Constitutive RELA over-expression drives increased expression of the panel of NF-κB-associated cytokines, which is suppressed by DOX addition (Figure 3I, Supplementary Figure 1B). Inhibition of POLE has a reduced to no effect on the expression of these cytokines both in the RELA constitutively over-expressed or knockout state, despite robust induction of DNA damage (Figure 3H-I, Supplementary Figure 1B). These data demonstrate that induction of inflammatory cytokines upon POLE induction is mediated by canonical NF-κB signaling mediated by RELA. In contrast, suppression of non-canonical NF-κB signaling via RELB suppression fails to substantially alter this transcriptional response (Supplementary Figure 1C-D).

A number of upstream pathways activate canonical NF-κB signaling depending on the cellular context and environmental stressor. To understand which of these pathways contribute to NF-κB activation upon POLE inhibition, we suppressed several of them individually and assessed the impact on the response to POLE inhibition. IKKγ is a protein implicated in activating canonical NF-κB signaling in response to both DNA damage and receptor mediated activation ^17,44^. Because we anticipated that loss of IKKγ would be detrimental to cell growth, we generated MDA-MB-231 cells expressing an sgRNA-resistant, codon optimized, DOX-repressible cDNA for *IKBKG* (encoding IKKγ) and disrupted endogenous *IKBKG* by CRISPR-Cas9 (DOX-OFF IKKγ). Addition of DOX reduced both phosphorylated and total levels of IKKγ (Supplementary Figure 1E) and blunted NF-κB cytokine induction following POLE suppression (Figure 3J).

We next considered the respective contributions of pattern recognition receptors (PRRs) activated by cytosolic DNA and RNA, known to be generated by replication stress. Cytosolic DNA can be recognized by cyclic GMP-AMP synthase (cGAS), which makes the second messenger cGAMP to activate stimulator of interferon genes (STING). Activation of the cGAS-STING pathway drives interferon and inflammatory signaling^45^. We disrupted STING using CRISPR/Cas9 in MDA-MB-231 cells and confirmed loss of the corresponding protein in single cell clones (Supplementary Fig 1F). Loss of STING results in a moderate blunting of inflammatory cytokine induction upon POLE suppression, without affecting the cytotoxic effects of POLE inhibition (Figure 3K, Supplementary Fig 1F). Retinoic Acid Inducible Gene I (RIG-I) functions as a PRR and responds by triggering robust innate immunity through a signaling cascade that stimulates downstream interferon transcription via both NF-κB-dependent and independent mechanisms^46^. We disrupted DDX58 (encoding RIG-I) in MDA-MB-231 cells using CRISPR/Cas9. Knockout was confirmed via immunoblot for RIG-I and assessing abrogation of RELA signaling in response to 3p-hpRNA, a RIG-I agonist (Supplementary Fig 1G-H). Upon POLE suppression, RIG-I knockout also moderately blunts NF-κB cytokine induction (Figure 3L). Taken together, these results demonstrate that canonical NF-κB signaling through RELA is required for the transcriptional upregulation of the NF-κB-associated cytokines identified in our transcriptomic analysis, and that this response is mediated in part through IKKγ, STING, and RIG-I, and reinforced by autocrine signaling (Figure 3M-N).

### POLE Inhibition Leads to Tumor Eradication and Robust Immune Infiltration

Given the robust NF-κB-mediated inflammatory signaling we observe *in vitro*, we next asked whether this inflammatory program could impact the tumor microenvironment (TME) and whether these effects would be immunostimulatory or immunosuppressive in the context of tumor growth. We used the DOXi shPOLE MDA-MD-231 model (Figure 3F) and generated orthotopic mammary fat pad tumors in athymic nude mice, which are largely deficient in adaptive immune responses. POLE inhibition induced marked tumor control, and upon withdrawal of DOX after 32 days (POLE reactivation), 5 of 6 tumors were unable to regrow (Figure 4A). We harvested tumors on Day 9 of DOX administration and observed significant infiltration and staining intensity of F4/80 positive cells (Figure 4B). These data indicate that POLE inhibition leads to robust tumor control and innate immune infiltration in xenograft models.

**FIGURE 4:**
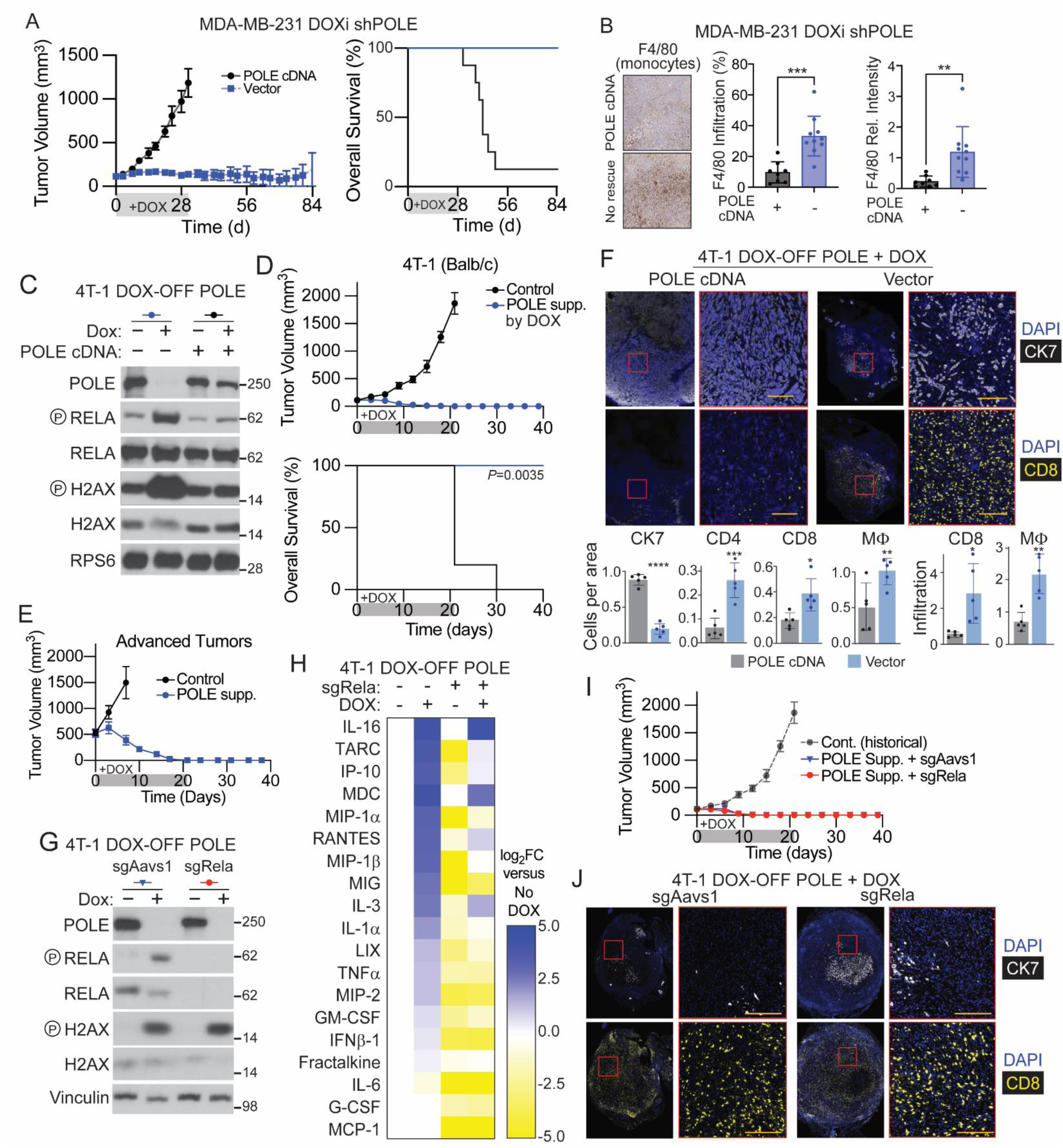
POLE Inhibition Leads to Tumor Eradication and Robust Immune Infiltration. **A,** Tumor volume (left, mm^3^) and survival (right) of NCr nude mice harboring orthotopic tumors derived from MDA-MB-231 cells stably expressing a DOX-inducible shRNA targeting POLE and either an shRNA-resistant POLE cDNA (black) or control vector (blue). DOX food added when tumors reached 100mm^3^ for 28d. **B,** Immunohistochemical stain (left) for F4/80 from tumors initiated as in A after 9d DOX treatment and quantification (right) of F4/80+ cell infiltration and intensity. **C,** Immunoblot for indicated proteins of lysates from a 4T-1 mouse mammary tumor cell clone expressing a DOX-repressible POLE cDNA and sgRNA targeting endogenous Pole. Cells were additionally transduced with a constitutive POLE cDNA and treated with DOX (100ng/mL, 3d), as indicated. **D-E,** Tumor volume (top, mm^3^) and survival (bottom) of Balb/c mice harboring orthotopic tumors derived from cells in ***C***. DOX food added when tumors reached 100mm^3^ (***D***) or 500mm^3^ (***E***) for 20d. **F,** Multiplex immunofluorescence (top) of tumors initiated as in ***D*** (9d DOX) for indicated markers and quantification (bottom) of positive cells per area or tumor infiltration for indicated populations, n=5 per group. **G,** Immunoblot for indicated proteins for cells from C transduced with an sgRNA targeting Rela (sgRela) or control (sgAAVS1) and treated with DOX (100ng/mL, 3d), as indicated. **H,** Fold change (log_2_) in the level of indicated cytokines and chemokines in cell culture media from the indicated cell lines from ***G***, relative to a cell line transduced with a POLE cDNA, n=2. **I,** Tumor volume (mm^3^) of Balb/c mice harboring orthotopic tumors derived from cells in ***G*** compared to a historical control (***D***). DOX food added when tumors reached 100mm^3^ for 20d. **J,** Multiplex immunofluorescence of tumors initiated as in ***I*** (9d DOX) for indicated markers, n=5 per group. Immunoblots are representative of n=3 independent replicates. Data report the mean and standard deviation, *** p<0.001, ** p<0.01, *p<0.05.

To suppress POLE more durably and in an immunocompetent host, we developed a near-complete POLE suppression TNBC allograft model in 4T1 mouse mammary tumor cells. We expressed a DOX-Repressible POLE cDNA in 4T1 cells with subsequent deletion of endogenous *Pole* (4T1 DOX-OFF POLE) and subsequent rescue with a constitutive POLE cDNA (Figure 4C). Upon POLE suppression, we observed loss of POLE protein and induction of H2AX phosphorylation (Figure 4C). POLE inhibition resulted in eradication of established orthotopic tumors within 12 days of DOX treatment, with no outgrowth observed after 20 days of DOX withdrawal (Figure 4D). Tumor eradication with similar kinetics was observed with reduced DOX treatment duration (3 or 6 days), low DOX concentration (2ug/mL), in large tumors (500 mm^3^, Supplementary Figure 2A-B and Fig 4E). Similar tumor eradication was observed in immunocompromised (NOD-Scid) mice (Supplementary Figure 2C), demonstrating that cancer cell-intrinsic responses upon near-complete POLE suppression are sufficient to induce tumor regression.

To understand the impact of POLE inhibition on the immune TME, we performed multiplexed immunofluorescence in DOX-OFF POLE inhibited tumors on day 9 of DOX addition. Consistent with tumor regression, total tissue area and staining for tumor marker CK7 were markedly reduced in POLE suppressed tumors (Figure 4F). The total number of F4/80+ macrophages, CD8+ and CD4+ T cells were significantly increased in POLE-suppressed tumors, as was infiltration of F4/80+ macrophages and CD8+ T cells (Figure 4F and Supplementary Figure 2D).

To gauge the contribution of RELA to immune infiltration in an immunocompetent allograft model, we expressed an sgRNA targeting Rela in DOX-OFF POLE cells and confirmed the absence of both phosphorylated and total levels of RELA protein. (Figure 4G). Similar to the results obtained in human TNBC cell lines, 4T1 cells exhibit an increase in the extracellular levels of 16 cytokines upon POLE suppression, 14 of which were substantially reduced by loss of RELA; and some cytokines that exhibited strong Rela dependence (IL-6, MCP1) were not induced by POLE loss (Figure 4H). RELA suppression does not prevent tumor regression upon POLE suppression, consistent with the observed cancer cell-intrinsic effects of POLE inhibition on tumor regression (Figure 4I). Multiplexed immunofluorescence revealed that immune cell infiltration was not impacted by RELA suppression (Figure 4J and Supplementary Figure 2E). These data demonstrate that POLE inhibition results in rapid tumor eradication via cancer cell-intrinsic mechanisms in an aggressive mouse TNBC model.

## DISCUSSION

In this study, we aim to understand the specific consequences of POLE suppression on cell signaling pathways and tumor biology. Our transcriptomic analysis reveals that partial POLE suppression in TNBC robustly activates a transcriptional program that does not overlap with partial POLD suppression or POLE suppression in LUBC cells. While genotoxic agents that directly target replication share transcriptional signatures with POLE suppression in TNBC, other genotoxic agents do not. Moreover, POLE suppression more robustly and selectively activates several genes, including multiple inflammatory cytokines, at the time points investigated. We find that the canonical NF-κB pathway is activated downstream of POLE suppression specifically in TNBC cells, resulting in the secretion of pro-inflammatory cytokines. As such, POLE suppression may offer a novel approach to engage anti-tumor immunity in TNBC.

Our work also demonstrates that POLE is an attractive anti-cancer target in TNBC through cancer cell-intrinsic mechanisms, as we observed dramatic and durable eradication of murine mammary tumors upon POLE inhibition. Supporting target selectivity, people harboring POLE mutations that reduce protein levels to less than 10% of wild-type are viable and survive to adulthood.^47^ Moreover, both fibroblasts from individuals with POLE mutations and non-TNBC cell lines in which POLE is partially suppressed, exhibit reduced rates of nucleotide incorporation, but otherwise have normal cell cycle progression, indicating that normal cells can tolerate low POLE levels that TNBC cells intrinsically cannot.^40,47^ TNBC frequently harbors alterations such as CCNE1 and MYC gain-of-function and p53 and RB loss-of-function that may abrogate cell cycle checkpoints needed to maintain viability in other breast cancer subtypes and non-transformed cells. Phosphoproteomic analysis of human breast cancer tumors reveals that CDK2 activity is elevated in TNBC compared to other breast cancer subtypes.^40^ These data provide a rationale for the selective vulnerability of TNBC to POLE inhibition, namely intrinsic activation of CDK2 downstream of TNBC genetic drivers and other factors.

Direct targeting of DNA replication may be less prone to common resistance mechanisms than widely-used nucleotide analogs. Many nucleotide analogs are prodrugs that require metabolic processes to produce toxic derivatives, resulting in well-established resistance mechanisms that block analog metabolism, increase catabolism, interfere with uptake, or upregulate *de novo* nucleotide synthesis.^48^ In contrast, replicative polymerases other than POLE are not sufficiently processive to facilitate genome duplication in human cells, and the requirement for POLE depends on its catalytic activity.^40^ Thus, resistance to POLE inhibition may be limited to general mechanisms including drug efflux, target mutation, or epigenetic changes, including quiescence or drug tolerant persister states that contribute to an immunosuppressive TME.^49,50^ Pharmacological approaches will be necessary to delineate the therapeutic efficacy of anti-cancer methods selectively targeting this vulnerability.

Immune system engagement is important to many successful cancer therapies. Indeed, tumor immune infiltration at diagnosis is a strong positive prognostic indicator in TNBC. Therefore, therapeutically targeting POLE may offer dual benefits. In addition to inducing tumor cell death, POLE suppression triggers NF-κB–mediated cytokine release, which may exert both autocrine and paracrine effects to amplify inflammatory signaling in neighboring cells. In this context, NF-κB activation could promote further cytokine release and the secretion of DAMPs, fostering a pro-inflammatory environment conducive to adaptive immune recruitment. Indeed, several cytokines and chemokines upregulated in POLE-suppressed MDA-MB-231 cells are known mediators of adaptive immune cell infiltration. While activation of inflammatory programs is not required to facilitate tumor eradication in the mouse TNBC model used, neither do they impede tumor clearance, as chronic NF-κB activation has been shown to enforce an immune-suppressive TME in certain contexts. Activation of replication stress has been proposed as a promising anti-cancer target in multiple contexts, including through inhibition of DNA repair and signaling pathways in TNBC such as POL θ, PARP, and the ATR/CHK1 pathway^36,51^. Given the recent success of immune checkpoint inhibitors in TNBC patients, understanding the interface between modulation of replication stress, inflammatory signaling, the TME, and immunotherapy response promises to be an active and fruitful area of future investigation in TNBC and other cancers.

Despite four decades of research on NF-κB signaling, the mechanisms by which it can be activated to trigger inflammatory signaling programs remain incompletely understood. Our data support a model in which inhibition of replication in TNBC distinctly triggers inflammatory signaling dependent on RELA and IKKγ, with contributions from DAMP receptors and reinforcement by cytokine-mediated autocrine and paracrine signaling. Apart from RELA, IKKγ may be particularly important for the response to replication stress due to its dual role in signaling pathways mediated by the DNA damage sensor ATM as well as receptor-mediated cytokine signaling.^17^

Our current understanding of NF-κB activation in breast cancer in response to therapy indicates that its impact on anti-tumor immunity and specific downstream signaling axes is also highly context dependent. NF-κB triggers the release of different immunomodulatory cytokines from cancer cells depending on tumor type and the class of genotoxic stress.^34,52,53^ For example, inhibition of ER signaling in LUBC supports NF-κB-mediated activation of interferon-stimulated genes, whose expression promotes tumor control.^14^ In TNBC, mitotic chromosomal instability activates NF-κB and interferon-stimulated genes mediated by RELB.^54^ Here, upon POLE inhibition we observe NF-κB activation mediated by RELA, resulting in a distinct pattern of target gene expression that favors pro-inflammatory chemo-attractants without interferon stimulation. These disparate examples of how NF-κB signaling contributes to inflammatory gene expression and tumor control in breast cancer suggest that additional work will continue to uncover a complex relationship between oncogenic pathways, genotoxic stress, NF-κB, and the TME.

## METHODS

### Reagents Used

Antibodies against Chk1 (sc-8408) from Santa Cruz; antibody against POLE (GTX132100) from GeneTex, antibodies against pH2AX S139 (9718), H2AX (2595), pChk1 S345 (2348), phospho p65 S536 (93H1), p65 (D14E12), phospho IKKy S376 (2689), phospho IKBA S32 (14D4), IKBA (44D4), IKKy (2685) and Anti-Mouse IgG (Alexa Fluor 488, 4410) from Cell Signaling Technologies, antibody against POLD (ab186407), and Goat Anti-Rat IgG H&L (Alexa Fluor 468, ab150160) from Abcam; MDA-MB-231, Hs578t, MCF7, 4T1 from ATCC; Matrigel (356234) and RPMI (10-040) from Corning; PrimeSTAR DNA polymerase (R040A) from Takara; pBabe-Puro-IKBalpha-mut (super repressor, #15291) and pBabe-Puro from Addgene; Doxycycline hyclate (AAJ6057914) from Fisher Scientific, human TNF-alpha recombinant protein (#16769) from Cell Signaling Technologies, Hydroxyurea (H8627), DAPI (D9542), 5-Bromo-2’-Deoxyuridine (B5002), and PhosSTOP (4906845001) from Sigma, MK-1775 (21266), Hybond-N+ membrane (RPN303B) from GE Healthcare, Quick Ligase (M2200) from New England Biolabs, Fetal Bovine Serum from Peak Serum, Pierce BCA Protein Assay (23225) from Fisher Scientific, RNeasy Plus Mini Kit (74136), Qiaprep Spin Miniprep Kit (27106), and QIAquick Gel Extraction Kit (28706) from Qiagen, Superscript IV (18090010) from Invitrogen, Polyethylenimine (239662) from Polysciences, Luciferase Assay System kit (E1500) from Promega.

Lentiviral shRNAs were obtained from the The RNAi Consortium (TRC) collection of the Broad Institute, or identical sequences cloned into the TRC vector (pLKO.1PS or Tet-pLKO-puro). The TRC website is: http://portals.broadinstitute.org/gpp/public/. The TRC#s for the shRNAs used are: shGFP, TRCN0000072186; shRFP, TRCN0000072203; shPOLE, TRCN0000052973; shPOLD1, TRCN0000352782.

sgRNAs were cloned into lentiCRISPRv2 (Addgene) linearized with BsmBI (NEB R0739L) using the following primers:

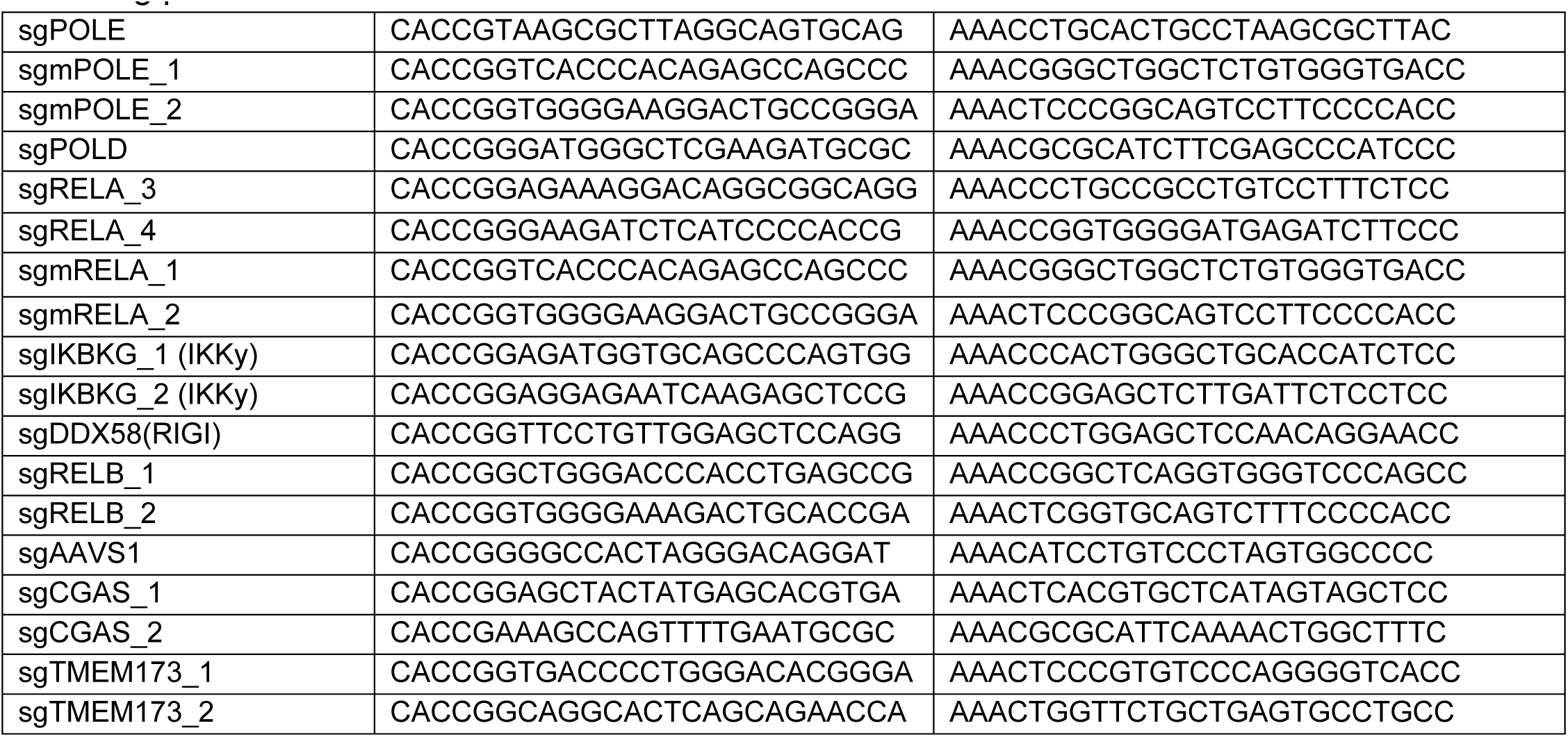

sgRNA-resistant versions of POLE WT, RELA and IKBKG cDNA was cloned into pCW57.1-MAT2A (Addgene 100521), replacing MAT2A. ShRNA resistant versions of POLE WT and POLE mutant cDNAs were also cloned into pMXS-IRES-Blast. Plasmids and expression vectors used in this study are deposited at Addgene (www.addgene.org) for distribution.

sgRNAs were cloned into lentiCRISPRv2 (Addgene) (TMEM, cGAS, DDX58, RELB, RELA) linearized with BsmBI (NEB R0739L) by Gibson assembly using NEBuilder HiFi DNA Assembly Master Mix (NEB E2621L).

### Animal Experiments

Tumors were initiated into 4-8 week old female CB17-Prkdc^scid^/J (NOD SCID, Jackson Labs), NCr athymic nude (Taconic), or Balb/c mice (Jackson Labs) orthotopically in the mouse mammary gland by implanting 500,000 cells in 33% matrigel into the 4^th^ murine mammary fat pad in a total volume of 25 μL. Expression of shPOLE_1 as well as repression of DOX-OFF POLE was induced by switching mice onto doxycycline chow (1000 mg/kg) upon formation of 100mm^3^ tumors. Tumor volume was assessed using caliper measurements and volume was calculated using (L*W^2^)/2. All experiments involving mice were carried out with approval from the Committee for Animal Care and under supervision of the Department of Comparative at NYU Langone Medical Center.

### Cell Culture

Cells were tested to be mycoplasma free by PCR based methods and authenticity verified by STR profiling (Duke University). Cells were cultured in RPMI supplemented with 10% IFS and penicillin/streptomycin.

### Generation of Recombinant Cell Lines

Lentiviral constructs were transfected with lentiviral packaging vectors ΔVPR and CMV VSV-G, while retroviral constructs were transfected with pCL-Ampho into HEK293 cells using polyethylenimine. Media was changed 12-16 hours post infection, and virus was collected 48 and 72 hours post infection and combined. Virus was passed through a 0.45 μm filter and stored at -80 °C or used immediately. One day prior to infection, cells were plated into 6-well tissue culture plates. Cells were infected with virus in media containing 1 μg/ml polybrene via spin infection in a Beckman Coulter Allegra X-12R centrifuge with an SX4750 rotor and Microplate Carrier attachment at 2,250 rpm for 30 min. The morning following infection cells were selected in puromycin for 3 days and then the media changed and cells allowed to recover for an additional day before being plated for experiments. Replating of cells for experiments occurred 5 days post infection.

### Generation of 231 DOX-OFF RELA and IKKγ models

Human MDA-MB-231 cancer cells were passaged in RPMI medium. One day prior to infection, cells were plated into a 6-well tissue culture plate. Cells were infected with virus expressing a *RELA* or *IKBKG* cDNA cloned into a pCW57.1 plasmid (Addgene Plasmid #100521) in media containing 1 μg/ml polybrene via spin infection in a Beckman Coulter Allegra X-12R centrifuge with an SX4750 rotor and Microplate Carrier attachment at 2,250 rpm for 30 min. A multiplicity of infection of 1 was used, and the morning after infection cells were selected with puromycin for 3 days and then the media was changed and cells were allowed to recover for an additional day before being plated for the subsequent rounds of infection. Following puromycin selection, cells were then plated for a second round of infection. They were infected with a virus expressing an sgRNA against *RELA* or *IKBKG* (cloned into the LentiCRISPRV2-hygro construct). The media was changed the following day and cells were allowed to recover for an additional day prior to changing the media to hygromycin for 7 days to induce selection. After 7 days of selection, the media was changed and cells were plated as single cells into 96-well plates in order to generate single cell clones. 3 days after plating for single cell clones, the plates were screened for the presence of single cell clones using a Zeiss Axiovert inverted microscope. Clones were grown for 10 days and then subjected to doxycycline addition and western blotting for RELA/IKKγ levels in order to determine successful generation of DOX-OFF RELA and DOX-OFF IKKγ clones.

### Generation of 4T1 DOX-OFF POLE models

Mouse mammary 4T1 cancer cells were passaged in RPMI medium. One day prior to infection, cells were plated into a 6-well tissue culture plate. Cells were infected with virus expressing a POLE cDNA cloned into a pCW57.1 plasmid (Addgene Plasmid #100521) in media containing 1 μg/ml polybrene via spin infection in a Beckman Coulter Allegra X-12R centrifuge with an SX4750 rotor and Microplate Carrier attachment at 2,250 rpm for 30 min. A multiplicity of infection of 1 was used, and the morning after infection cells were selected with puromycin for 3 days and then the media was changed and cells were allowed to recover for an additional day before being plated for the subsequent rounds of infection. Cells were replated and subsequently infected with a virus containing a POLE cDNA cloned into an MXS vector utilizing the same conditions as the previous infection. The morning after infection, cells were selected with blasticidin for 5 days. Following recovery from blasticidin selection, cells were then plated for a third round of infection. They were infected with a virus expressing an sgRNA against *Pole1* (cloned into the LentiCRISPRV2-hygro construct). The media was changed the following day and cells were allowed to recover for an additional day prior to changing the media to hygromycin for 7 days to induce selection. After 7 days of selection, the media was changed and cells were plated as single cells into 96-well plates in order to generate single cell clones. 3 days after plating for single cell clones, the plates were screened for the presence of single cell clones using a Zeiss Axiovert inverted microscope. Clones were grown for 10 days and then subjected to doxycycline addition and western blotting for POLE levels in order to determine successful generation of DOX-OFF POLE clones.

### Proliferation Assays

Direct cell counts were performed using a Beckman Z2 Coulter Counter with a size selection setting of 8 to 30 μm. Population doublings were calculated based on the ratio of cells at end of experiment to cell counts at time of withdrawal. For all other proliferation assays, 25,000 cells were plated in triplicate into 6-well plates for 5 days.

### Immunoblotting

Lysates collected on ice by washing cells in cold PBS followed by addition of lysis buffer (50 mM Tris pH 7.4, 150 mM NaCl, 1% NP-40, 0.1% sodium deoxycholate, 0.1% SDS, 2 mM EDTA) containing a protease inhibitor cocktail (Roche) and PhosSTOP. Lysates were then incubated on ice for 10 min and sonicated for 10s per sample in a cold room. Protein levels were quantified using a BCA protein assay kit (Pierce) and 8 µg protein was loaded and electrophoresed onto Bolt 4–12% Bis–Tris polyacrylamide gels (Thermo Fisher) or 3-8% Tris-Acetate polyacrylamide gels (Thermo Fisher) for POLE detection. Gels were transferred onto a PVDF membrane (Millipore IPVH00010) in transfer buffer (2.2 g/L CAPS, 0.45 g/L NaOH, 10% ethanol) for 2 hours at 60V for Bis-Tris gels and 3 hours at 60V for Tris-Acetate gels.

### Correlations of Publicly Available Data

Sensitivity scores were downloaded from https://depmap.org/portal/download/all/. File Name: ExpandedGeneZSolsCleaned.CSV. To obtain a ranked list of gene expression correlations to POLE suppression in breast cancer cell lines, CCLE expression data and sensitivity score data were filtered for breast cancer cell lines and a Pearson correlation was obtained between sensitivity score and gene expression. GSEA was performed using the ranked list with 1000 permutations and the curated gene sets.

### DepMap analysis

Differential genetic dependency analysis between TNBC and non-TNBC cell lines was derived from the RNAi gene dependency dataset (Achilles+DRIVE+Marcotte, DEMETER2^55^), consisting of 16,810 genes, utilizing the two class comparison tool on the depmap portal (www.depmap.org). Cancer cell lines classified as TNBC (N=31): BT20, BT549, CAL120, CAL148, CAL51, CAL851, DU4475, HCC1143, HCC1187, HCC1395, HCC1599, HCC1806, HCC1937, HCC2157, HCC2185, HCC3153, HCC38, HCC70, HDQP1, HS578T, MDAMB157, MDAMB231, MDAMB436, MDAMB468, MFM223, SKBR7, SUM102PT, SUM1315MO2, SUM149PT, SUM159PT, SUM185PE, SUM229PE^56^. All other breast cancer cell lines were classified as non-TNBC (N=49).^57^

### qPCR

RNA was isolated by column purification (RNeasy Kit, Qiagen) and cDNA synthesis was performed by reverse transcription of 1 µg of total RNA by reverse transcriptase (Superscript IV, 18090010, Invitrogen) in a reaction containing 1 µl RNase OUT (10777019, Invitrogen). qPCR was performed on cDNA using SYBR green quantification (Maxima qPCR master mix, K0222, Thermo Fisher). All genes were quantified relative to RPL13A. The following primers used:

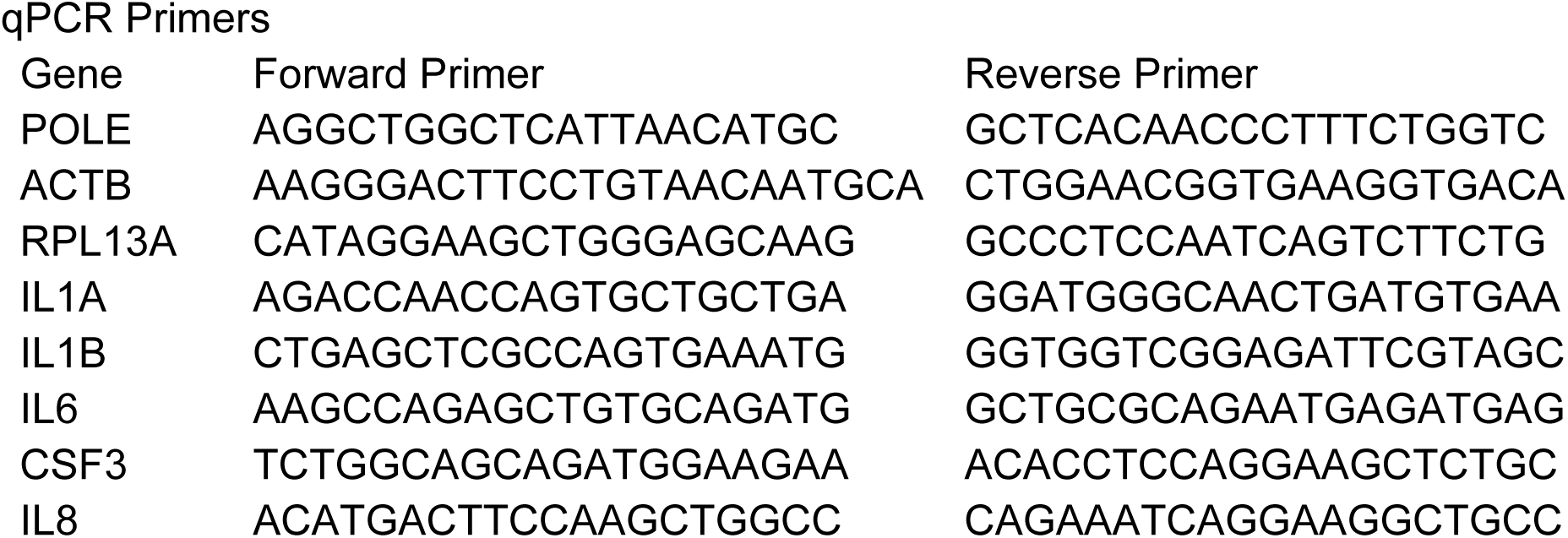

### RNAseq

MDA-MB-231, HS578t, and MCF7 cells stably transduced with an shRNA-resistant POLE or POLD cDNA (in pMXS-blast) in media containing 1 μg/ml polybrene via spin infection in a Beckman Coulter Allegra X-12R centrifuge with an SX4750 rotor and Microplate Carrier attachment at 2,250 rpm for 30 min. A multiplicity of infection of 1 was used, and the morning after infection cells were selected with blasticidin for 3 days and then the media was changed and cells were allowed to recover before being plated for the subsequent rounds of infection. Cells were replated and subsequently infected with shPOLE or shPOLD virus using the conditions described above. A multiplicity of infection of 1 was used, and the morning after infection cells were selected with puromycin for 5 days of selection. Following 5 days of selection, the media was changed, and cells were collected for subsequent RNA isolation. Parent MDA-MB-231 cells were also treated with hydroxyurea (2mM), aphidicolin (2ug/ml), paclitaxel (4nM), cycloheximide (2uM) and cisplatin (12uM) and DMSO for 24 hours and cells were collected for subsequent RNA isolation. RNA was isolated using the RNeasy Plus Mini Kit. Analysis was done with normalized read counts. GSEA was performed with 1000 permutations in the curated gene sets.

### Conditioned Media

Indicated cell lines were plated in a 6-well tissue culture plate 6-well plate and infected with shGFP and shPOLE per protocol for 5 days. On day 4 of treatment, media was changed to fresh media with 1ml of media per well and parent cell lines were plated. On day 5, conditioned media was collected, spun down at 1200g for 5 min to remove floating cells/debris and added to the plated parental cell lines for indicated time points. Following incubation, cells were collected for further RNA isolation.

### Cytokine Release

Indicated cell lines were plated in a 6-well plate and either infected with shGFP and shPOLE as above for 5 days or treated with 0.5ug/ml doxycycline for 5 days. On day 4 of treatment, media was changed to fresh media with 1ml of media per well. On day 5, media was collected, spun down at 1200g for 5 min to remove floating cells/debris, aliquoted into 100ul tubes. Samples were shipped to Eve Technologies for analysis.

Arrays used:

Human Cytokine/Chemokine Panel A 48-Plex Discovery Assay® Array (HD48A)

The Mouse Cytokine/Chemokine 44-Plex Discovery Assay® Array (MD44)

### Multiplexed Immunofluorescence

After fixing in formalin for 24 hours, tissues were dehydrated through graded ethanols and xylene and infiltrated with paraffin (Paraplast X-tra, Leica SKU 39603002) on a Leica Peloris II tissue processor. Embedded tissues were sectioned at 5 µm and stained with hematoxylin (Leica, 3801575) and eosin (Leica, 3801619) on a Leica ST5020 automated stainer to evaluate histology. Adjacent sections were immunostained on a Leica BondRx autostainer, according to the manufacturers’ instructions. In brief, sections were first treated with peroxide to inhibit endogenous peroxidases followed by an antigen retrieval step with ER2 (Leica 9AR9640) for 20 minutes at 100o. For standard immunohistochemistry, sections were incubated with the F4/80 antibody against macrophages (CST, Cat# 70076S) at a 1:1000 dilution for 30 minutes at room temperature. Primary rabbit antibodies were detected with anti-rabbit HRP-conjugated polymer and 3,3’-diaminobenzidine (DAB) substrate that are provided in the Leica BOND Polymer Refine Detection System (Cat # DS9800). For multiplex immunofluorescence staining with Akoya Biosciences® Opal™ reagents, slides were incubated with the first primary antibody and secondary polymer pair and then underwent HRP-mediated tyramide signal amplification with a specific Opal® fluorophore. The primary and secondary antibodies were subsequently removed with a heat retrieval step, leaving the Opal fluorophore covalently linked to the antigen. This sequence was repeated with subsequent primary and secondary antibody pairs and a different Opal fluorophore at each step (see table below for reagent details; the 4plex panel was identical to the 6plex panel without the Ki67 and Foxp3 antibodies). Sections were counterstained with spectral DAPI (Akoya Biosciences, FP1490) and mounted with ProLong Gold Antifade (ThermoFisher Scientific, P36935). Semi-automated image acquisition was performed on a Akoya Vectra Polaris (PhenoImagerHT) multispectral imaging system at 20X magnification using PhenoImagerHT 2.0 software in conjunction with Phenochart 2.0 and InForm 3.0 to generate unmixed whole slide qptiff scans.

| Antigen | Clone | Vendor, Cat# | RRID | Dilution Factor | Incubation Time | Secondary/<br>Polymer | Opal TSA fluorophore, | Fluorophore dilution |
| --- | --- | --- | --- | --- | --- | --- | --- | --- |
| F480 | D2S9R | CST, 70076S | AB_2799771 | 1500 | 60 | Rat- HRP polymer 1-Step (Biocare, BRR4016H) | 480 | 200 |
| CD4 | D7D2Z | CST, 25229S | AB_2798898 | 300 | 30 | Rabbit-on-Rodent HRP polymer (Biocare RMR622) | 620 | 150 |
| FoxP3 | D6O8R | CST, 12653S | AB_2797979 | 4000 | 30 | Rabbit-on-Rodent HRP polymer (Biocare RMR622) | 520 | 150 |
| CD8 | D4W2Z | CST, 98941 | AB_2756376 | 300 | 30 | Rabbit-on-Rodent HRP polymer (Biocare RMR622) | E570 | 150 |
| Ki67 | Sp6 | Abcam, Ab16667 | AB_302459 | 150 | 30 | Rabbit-on-Rodent HRP polymer<br>(Biocare<br>RMR622) | 690 | 150 |
| CK7 | EPR170<br>78 | Abcam,<br>ab181598 | AB_27838<br>22 | 4000 | 60 | Rabbit-on-Rodent<br>HRP<br>polymer<br>(Biocare<br>RMR622) | 780_TSA-DIG | 25_100 |

### Multiplexed Immunofluorescence Analysis

Immunofluorescence images were first analyzed by performing Watershed nucleus detection on the DAPI channel using QuPath^58^. Identification of tissue boundaries were performed using a Random Forest classifier for pixels provided through ilastik^59^. Subsequent analysis of cell frequencies and spatial localization were performed with MATLAB/R2022b (Mathworks). Gating thresholds were first manually set based on fluorescent intensities of the following stains: F4/80, CD4, CD8, Foxp3, Ki67, and CK7. For each nuclei detected, cell type was determined based on these gates. Furthermore, the image mask representing tumor tissue was divided into a central region and a peripheral region. The peripheral region is defined as the area within 40 μm of the tissue margins. For each cell type, the following 2 metrics were then quantified: (1) the number of cells per unit of pixel area of tissue within the central region and (2) the ratio of this cell density metric within the central region relative to the peripheral region. The latter provides information regarding the degree of cell infiltration deep into the tumor parenchyma.

### Immunohistochemistry Analysis

For quantification of F4/80 immunoreactivity within tumor sections, images were first down-sampled to a resolution of 1000 DPI (dots per inch). The ilastik pixel classifier^58^ was used to: (1) identify F4/80 immunoreactive puncta, (2) create an image mask representing central regions of tumor tissue, and (3) create an image mask representing the tissue margin, which was excluded from analysis. We calculated the degree of F4/80 infiltration by the following 2 means: (1) density quantification of F4/80 puncta density per unit pixel area (of the tumor central region) and (2) the inverse of the average of the Euclidean RGB distance from R = 0.2, G = 0.1, B = 0.1 (on a 0 to 1 scale) for each pixel in the central tumor region. This latter metric is indicative of the average intensity of immunoreactivity.

### Quantification and Statistical Analysis

For quantitative data, data was presented as mean ± SEM a two-tailed t-test was used for statistical analysis using Prism software for comparison of normal distribution samples. For multiple comparisons, a one-way analysis of variance (ANOVA) or multiple t-test was used as specified for each experiment in the figure legend with P values calculated in Prism. A p value less than 0.05 is statistically significant. Other statistical parameters such as number of experiment repeats and n values can be found in the figure legends. Studies were not conducted blind.

## ACKNOWLEDGEMENTS

We thank Possemato lab members for helpful comments. plentiCRISPR v2 was a gift from Feng Zhang, pBabe puro and pBabe-Puro-IKBalpha-mut were gifts of William Hahn. **Funding:** National Institutes of Health R01CA214948 (RP), R01GM132491 (RP), R01CA286141 (RP), F30CA271623 (EFS), American Cancer Society (RP), Cancer Center Support Grant P30CA016087 (Neel). **Author contributions:** Conceptualization: EFS, KMF, RP. Methodology: EFS, KMF, CL, JD, KKW, RP. Investigation: EFS, KMF, AT, DE, VOS, HM, TRK, JD; Writing: EFS, KMF, RP. Funding acquisition, resources, and supervision: RP.

---

**SUPPLEMENTARY FIGURE 1:**
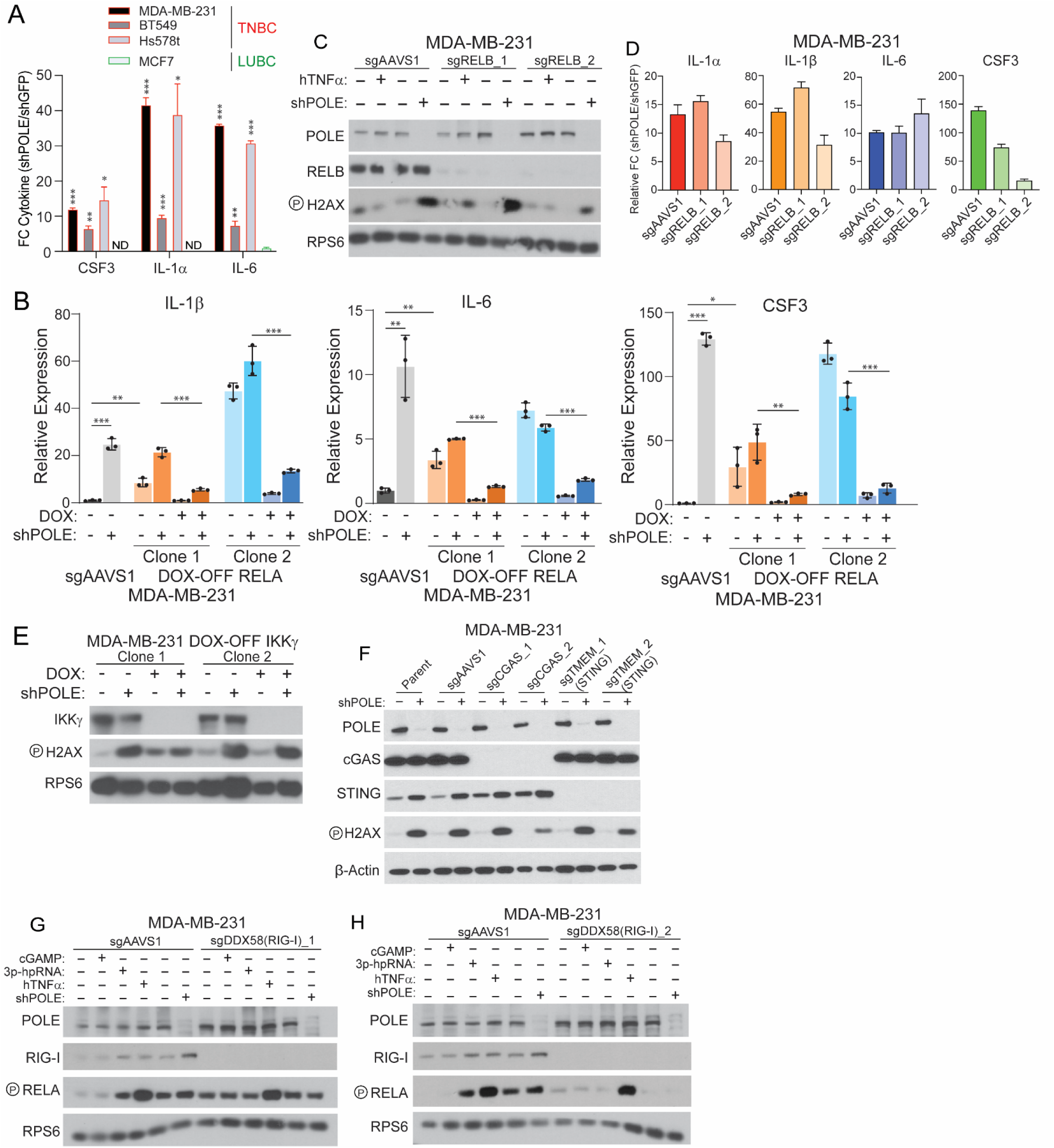
Data Supporting Figure 3. **A,** Fold change (log_2_) in the level of indicated cytokines and chemokines in cell culture media from the indicated cell lines infected with an shRNA targeting POLE, relative to a control shRNA (shGFP), 5d. **B,** mRNA level of the indicate gene in cell lines (Fig. 3H) or those expressing a control sgRNA (sgAAVS1), relative to sgAAVS1 clones transduced with shGFP (5d). **C,** Immunoblots for indicated proteins of lysates from MDA-MB-231 cell clones expressing one of two sgRNAs targeting RELB (RELB_1, RELB_2) or a control (sgAAVS1) with additional transduction of an shRNA targeting POLE (shPOLE, “+”) or a control (shGFP, “-“), 5d. **D,** mRNA level of indicated genes from MDA-MB-231 knockout clones transduced with shPOLE or shGFP from ***C***, relative to shGFP transduced cells in each condition. E, Immunoblots for indicated proteins of lysates from MDA-MB-231 cell clones expressing a DOX-repressible IKKγ cDNA and endogenous *IKKγ* disruption with additional transduction of an shRNA targeting POLE (shPOLE, “+”) or a control (shGFP, “-“) and DOX treatment (100ng/mL), as indicated (5d). F-H, Immunoblots for indicated proteins of lysates from MDA-MB-231 cell clones expressing sgRNAs targeting the indicated gene and additional transduction with an shRNA targeting POLE (shPOLE, “+”) or control (shGFP, “-“) for 5d and treatment with 2,3-cGAMP (5 uM for 30 minutes), 3p-hpRNA (1ug/ml for 30 minutes), or hTNFα (20ng/ml for 30 minutes) for 1d, as indicated. Immunoblots are representative of n=3 independent replicates. Data in ***A***, ***B***, ***D***, report the mean and standard deviation of n=3 biological replicates. *** p<0.001, ** p<0.01, *p<0.05.

**SUPPLEMENTARY FIGURE 2:**
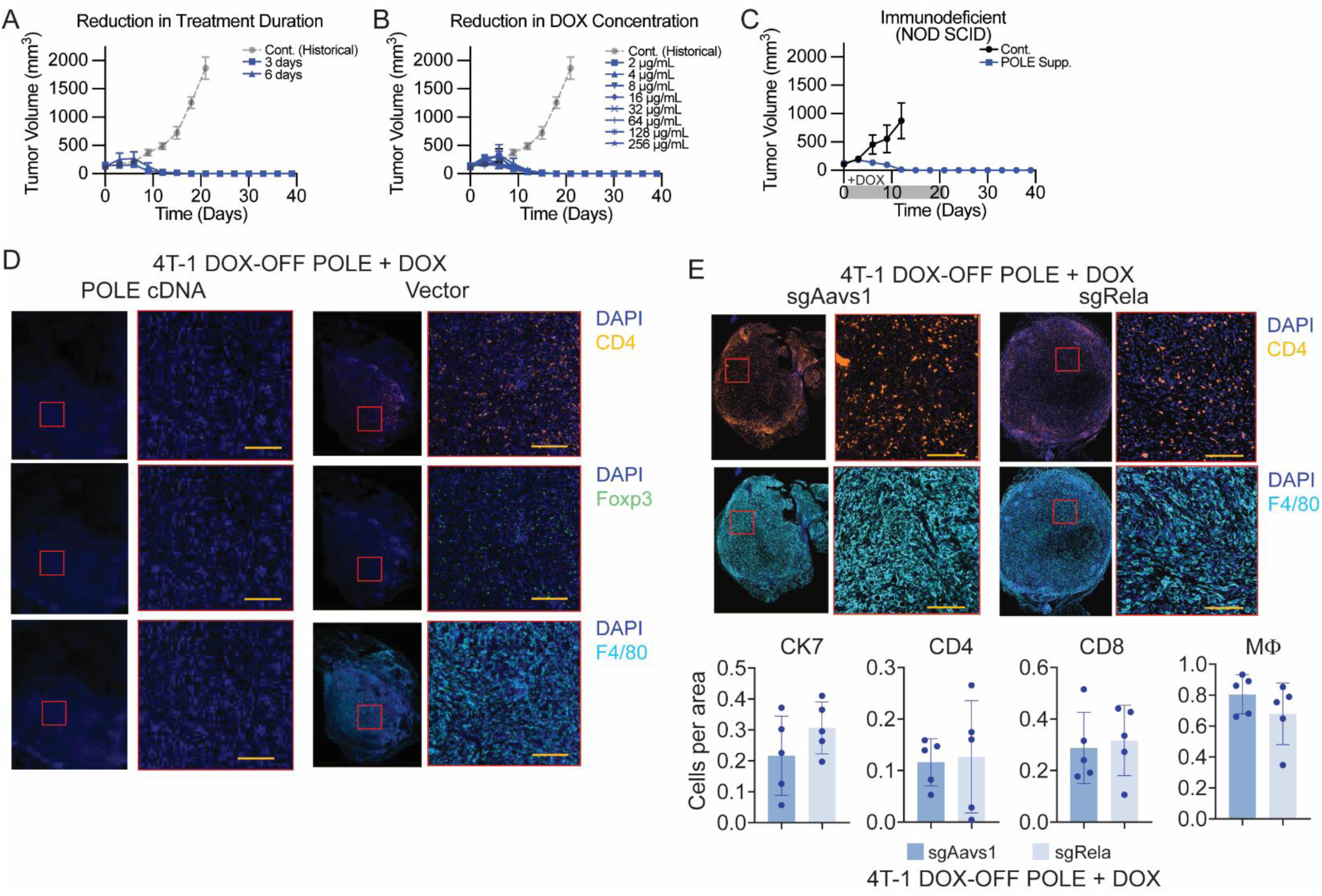
Data Supporting Figure 4. **A-C,** Tumor volume (mm^3^) of Balb/c (**A-B**) or CB17-Prkdc^scid^/J (NOD SCID, **C**) mice harboring orthotopic tumors derived from a 4T-1 mouse mammary tumor cell clone expressing a DOX-repressible POLE cDNA and sgRNA targeting endogenous Pole and additionally transduced with a constitutive POLE cDNA or control vector, as indicated (as in Fig. 4C). Mice were treated with DOX chow for indicated duration (***A*** and ***C***) or DOX water at indicated concentration continuously (***B***), relative to a historical control (Fig. 4D). **D,** Multiplex immunofluorescence of tumors initiated as in Fig. 4D (9d DOX) for indicated markers, n=5 per group. **E,** Multiplex immunofluorescence (top) of tumors initiated as in Fig. 4I (9d DOX) for indicated markers and quantification (bottom) of positive cells per area for indicated populations, n=5 per group. Data report the mean and standard deviation.

